# Cyclophilins A and B Oppositely Regulate Renal Tubular Epithelial Phenotype

**DOI:** 10.1101/288886

**Authors:** Eduard Sarró, Mónica Durán, Ana Rico, Anthony J. Croatt, Karl A. Nath, Salcedo Maria Teresa, Justin H. Gundelach, Daniel Batlle, Richard J. Bram, Anna Meseguer

## Abstract

Cyclophilins (Cyp) are peptidil-prolyl-isomerases and the intracellular receptors for the immunosuppressant Cyclosporine-A (CsA), which produces epithelial-mesenchymal-transition (EMT) and renal tubule-interstitial fibrosis. Since CsA inhibits Cyp enzymatic activity, we hypothesized that Cyp could be involved in EMT and fibrosis. Here, we demonstrate that CypB is a critical regulator of tubule epithelial cell plasticity on the basis that: i) CypB silencing caused epithelial differentiation in proximal tubule-derived HK-2 cells, ii) CypB silencing prevented TGFβ-induced EMT in HK-2, and iii) CypB knockdown mice exhibited reduced UUO-induced inflammation and kidney fibrosis. By contrast, silencing of CypA induces a more undifferentiated phenotype and favors TGFβ effects. EMT mediators Slug and Snail were up-regulated in CypA-silenced cells, while in CypB silencing, Slug, but not Snail, was down-regulated; thus, reinforcing the role of Slug in kidney fibrosis. CypA regulates Slug through its PPIase activity whereas CypB depends on its ER location, where interacts with calreticulin, a calcium modulator which is involved in TGFβ signaling. In conclusion, this work uncovers new roles for CypA and CypB in modulating proximal tubular cell plasticity.

## Introduction

Kidney fibrosis is the principal process underlying the progression of chronic kidney disease (CKD) to end stage renal disease (ESRD). Since specific therapies to prevent, slow down, or reverse fibrosis are severely lacking (1), understanding the complex molecular mechanisms and cellular mediators of kidney fibrosis could offer new therapeutic avenues to prevent the loss of kidney function (2).

Maladaptive repair due to repeated or sustained injury to the proximal tubule epithelial cells (PTC) has proved to be sufficient to induce CKD and fibrosis (3). Injured PTC, which drive injury and inflammation by releasing pro-fibrotic factors, in particular TGFβ (4), and by producing inflammatory cytokines, including TNFα (5, 6), have been identified as a major player in fibrosis. A very marked and relevant feature of kidney fibrosis is the transition of tubular epithelial cells into cells with mesenchymal features, a so-called epithelial-mesenchymal transition (EMT) (7). This switch in cell differentiation and behavior is mediated by transcription factors, including Snail (Snail1), Slug (Snail2), zinc-finger E-box-binding homeobox (ZEB)1/2 and Twist1/2, which negatively regulate E-cadherin expression through their recognition of common E-box sequences in the E-cadherin promoter (8, 9). Down-regulation of E-cadherin is the hallmark of EMT to reinforce the destabilization of adherens junctions in the epithelial barrier. In vivo, it has been proved that Snail is responsible for the partial EMT program (EMT2) that leads to dedifferentiation of renal epithelial cells and promotes kidney fibrosis (10, 11). Damaged epithelial cells undergoing EMT2 remain confined to the tissue without engaging in the delamination and invasion programs occurring in cancer (EMT3) (11).

The potent immunosuppressant Cyclosporine A (CsA) produces severe renal tubule-interstitial fibrosis that limit the drug’s clinical use (12). In vitro, CsA treatment of PTC induces a dose-dependent release of TGF-β, EMT events and increased expression of Snail (13), as well as, production of pro-inflammatory cytokines (14). Intracellularly, CsA binds to cyclophilins, a family of ubiquitous and highly conserved proteins that accelerate protein folding by catalyzing the cis-trans isomerization of proline residue (15, 16). CsA binding to cyclophilins inhibits their peptidyl-prolyl cis-trans isomerase (PPIase) activity. Cyclophilins do also play a prominent role as chaperones by mediating protein trafficking, protein-protein interactions, and as scaffolding proteins for assembly of macromolecular complexes (17). In humans, cyclophilins A (CypA) and B (CypB), the best characterized members of the family, are located in the cytosol and in the endoplasmic reticulum (ER), respectively (17). CypB controls ER homeostasis by participating in the protein quality control process in the ER (18) and is released to the extracellular media in the presence of CsA (19). Moreover, CypA and CypB can also be released in response to inflammatory stimuli and elevated circulating levels for both of them have been reported in different inflammatory diseases (20–22). They contribute to the inflammatory responses via their potent chemotactic properties for various immune cells (23, 24), which is mediated by the signaling receptor CD147 on target cells (25). In the kidney, CypB was found to be the interacting partner of KAP, a protein exclusively expressed in proximal tubule cells of the kidney that protects against CsA toxicity (26, 27), as well as interacting with sodium-potassium ATPase, being required for pump activity in proximal tubule cells of the kidney (28).

Considering the loss of epithelial phenotype triggered by CsA and its inhibitory actions on cyclophilins, the idea that the latest could be critical in the development of EMT and kidney fibrosis is entirely plausible. To investigate the potential involvement of cyclophilins in the regulation of the epithelial phenotype, we studied the effects of CypA and CypB silencing in human proximal tubule epithelial cells. We observed that CypA and CypB silencing exert opposite effects on the phenotype of proximal tubular cells by promoting or haltering, respectively, EMT processes through distinctly modulating the Snail family of epithelial repressors. Moreover, we also showed a marked attenuation of the molecular changes associated with inflammation and fibrosis in the kidneys of CypB KO mice subjected to unilateral ureteral obstruction (UUO). Results from this study pinpoint CypB as a potential therapeutic target to prevent fibrosis.

## Results

### CypB and CypA silencing differentially affects epithelial phenotype of cultured PTC

To investigate the potential involvement of cyclophilins in the regulation of the epithelial phenotype, we silenced CypA and CypB in HK-2 cells, a widely characterized proximal tubular epithelial cell line retaining a phenotype indicative of well-differentiated PTC (Fig. 1A). Since PTC in culture progressively acquire epithelial features upon reaching confluence, we first analyzed the expression levels of the epithelial markers E-cadherin (adherens junctions), ZO-1 and occludin (tight junctions) and keratin (intermediate filaments) at 2, 5 and 10 days post seeding, considering that cells reach confluence by the second day. We observed that all epithelial markers increased along days of culture (Fig. 1B). Five days post seeding was selected for further experiments. Our results show that CypB silencing greatly increased E-cadherin and occludin expression and to a lesser extent ZO-1 and keratin (Fig. 1C). By contrast CypA silencing reduced occludin, ZO-1 and keratin levels. Those results were also observed at the mRNA level (Fig. 1D). As shown in Figure 1E, the increase in E-cadherin expression observed in CypB-silenced cells correlated with augmented E-cadherin levels in the plasma membrane, suggesting a concomitant gain in E-cadherin functionality. Since a stimulatory role for E-cadherin in proliferation has been described (29), we explored the effect of CypB knockdown on HK-2 proliferation. In accordance with E-cadherin levels, cells lacking CypB showed higher proliferation indices than control cells, whereas CypA silencing had no significant effect (Fig 1F). By contrast, we observed that CypA silencing reduced transepithelial electric resistance (TER) and increased FITC-labeled Dextran permeability (Fig. 1G and 1H, respectively), which correlated with the downregulation in ZO-1 and occludin levels observed in CypA-silenced cells.

**Figure 1.**
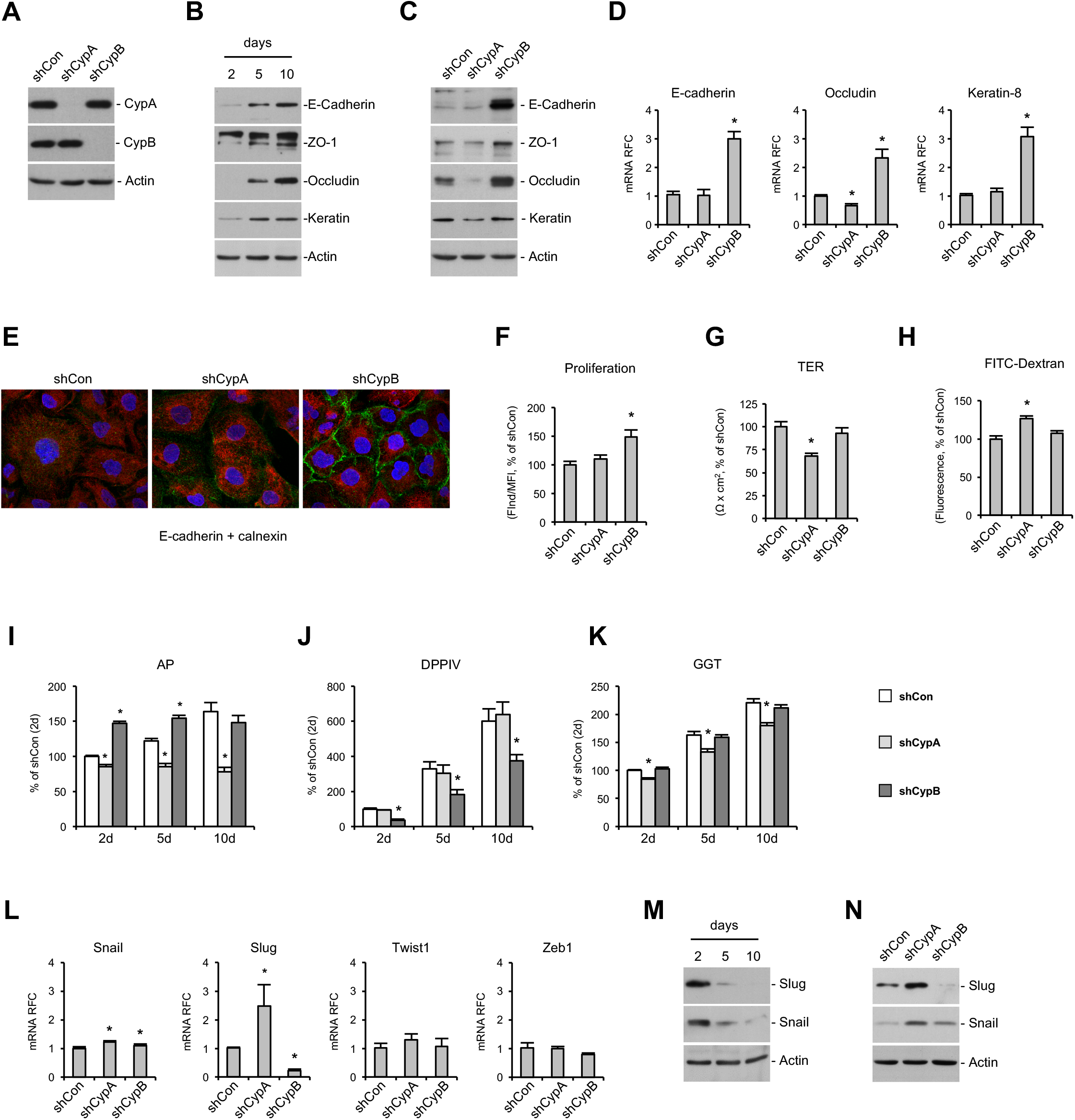
CypB and CypA silencing differentially affects epithelial phenotype of cultured PTC. To investigate the potential involvement of cyclophilins in the regulation of the epithelial phenotype, CypA and CypB were silenced in the proximal tubular epithelial cell line HK-2. (A) Western Blot showing the decrease in CypA and CypB expression in HK-2 cells stably transfected with shRNA-expressing lentiviral vectors against CypA or CypB, respectively. (B) The expression levels of the epithelial markers E-cadherin, ZO-1, occludin and keratin in HK-2 wild-type cells after 2, 5 and 10 days of culture were analyzed by western blot. The expression levels of the above epithelial markers in CypA and CypB-silenced cells after 5 days of culture were determined by immunoblotting (C) and real time quantitative PCR (qPCR) (D). (E) Immunofluorescence staining of E-cadherin (green) in CypA and CypB-silenced cells cultured for 5 days, showing membrane location of E-cadherin in CypB-silenced cells. Calnexin (red) was used to stain endoplasmatic reticulum. (F) Cell proliferation in control and CypA and CypB-silenced cells was measured by means of carboxyfluorescein succinimidyl ester (CFSE) labeling followed by flow cytometry analysis as indicated in Methods. Values indicate the ratio FInd/MFI, where MFI was the median fluorescence intensity of all viable cells at collection and Find the peak fluorescence intensity of the viable non-divided cells. To assess monolayer integrity after CypA and CypB-silencing, transepithelial electric resistance (TEER) (G) and FITC-labeled Dextran permeability (H) were measured in cyclophilin-silenced HK-2 cells cultured for 5 days. Enzymatic activities of alkaline phosphatase (AP) (I), dipeptidyl peptidase-IV (DPP-IV) (J) and gamma-glutamyltransferase (GGT) (K) in CypA and CypB-silenced cells after 2, 5 and 10 days of culture. (L) The expression levels of the transcriptional repressors Snail, Slug, Twist1 and Zeb1 were analyzed by real time quantitative PCR (qPCR). Finally, levels of Slug and Snail in HK-2 wild-type cells after 2, 5 and 10 days of culture (M) or in CypA and CypB-silenced cells at 5 days of culture (N) were analyzed by Western Blot. Student’s t-test was used to compare shCon vs shCypA or shCon vs shCypB for each culture time point. * P<0.05.

To further characterize the effects of cyclophilin silencing in epithelial differentiation, we analyzed the activities of the proximal tubule brush border enzymes alkaline phosphatase (AP), dipeptidyl peptidase-IV (DPP-IV) and gamma-glutamyltransferase (GGT) as recognized markers of epithelial differentiation. Aligned with the above results, AP activity was markedly enhanced in cells lacking CypB, with maximal levels already achieved after two days of culture (Fig. 1I). By contrast, DPPIV activity stayed below control levels (Fig. 1J) and GGT activity behaved as in controls (Fig. 1K). Diminished DPPIV activity might not be detrimental to the renal epithelial cells phenotype since it has been reported that DPPIV inhibitors block TGFβ signaling, thus protecting against renal fibrosis (30, 31). In CypA-silenced cells AP activity remained permanently low (Fig. 1I), DPPIV activity behaved as in control cells (Fig. 1J) and GGT activity increased over time but remained below control levels (Fig. 1K).

The temporal expression pattern of epithelial markers is tightly regulated by epithelial repressors, including the transcription factors Snail (Snail1), Slug (Snail2), zinc-finger E-box-binding homeobox (ZEB)1/2, and Twist1/2 (8, 9). Our results show that in HK-2 cells CypA silencing increased Slug mRNA levels and to a lesser extent those of Snail (Fig. 1L). Interestingly, CypB silencing also slightly increased the mRNA levels of Snail, but strongly reduced those of Slug (Fig. 1L). By contrast, neither CypA nor CypB silencing had any significant effect on Twist1 and Zeb1 mRNA levels (Fig. 1L). To corroborate these results, we analyzed the protein levels of Slug and Snail. Our results show that Slug and Snail expression progressively decreased from day 2 to 10 (Fig. 1M), correlating with the augmented epithelial marker expression observed in Figure 1B. Moreover, at 5 days post-seeding, we observed that Slug protein levels were increased in CypA-silenced cells while they were undetectable in CypB-silenced cells (Fig. 1N). Both CypA and CypB increased Snail protein levels to a higher extent than that observed at the mRNA level, suggesting additional regulatory mechanisms besides transcriptional modulation (Fig. 1N).

In order to corroborate these results, we silenced CypB and CypA in another proximal tubule derived cell line (RPTEC) (Fig S1). Our results show that, as in HK-2 cells, RPTEC cells lacking CypB presented reduced slug expression and higher Snail levels. E-cadherin upregulation in CypB silenced cells was less appreciable in RPTEC cells, mostly due to the higher basal expression of this protein in comparison with HK-2 cells.

Taken together, these results indicate that while CypB is necessary for Slug expression, CypA acts as a repressor of both Slug and Snail. Moreover, our results also indicate that, at least in CypB-silenced cells, loss of Slug rather than gain of Snail prevail in the regulation of epithelial markers.

### Cyclophilin A and B exert divergent effects on TGFβ action on epithelial cells

TGFβ has been widely regarded as a primary factor driving renal fibrosis (4). In vitro, TGFβ treatment alone can induce proximal tubule cells to undergo an epithelial-to-mesenchymal transition (EMT) (4, 32). Considering the results shown in Figure 1, we decided to investigate the effects of CypB and CypA silencing on TGFβ signaling. In HK-2 cells, TGFβ induced an EMT-like process demonstrated by a gradual decrease of E-cadherin and occludin expression and an increase of Fibronectin levels that were not related with changes in CypA or CypB levels (Fig. 2A). Reminiscent of what happens in basal conditions, it is of note that CypB silencing partially prevented TGFβ-induced EMT by maintaining E-cadherin and occludin levels closer to control and reducing fibronectin expression (Fig. 2B). In addition, CypA silencing enhanced TGFβ-induced EMT by further decreasing E-cadherin and occludin levels and increasing those of fibronectin. Loss of E-cadherin and occludin expression after TGFβ treatment was consistent with a switch from a well defined monolayer with cobblestone morphology of control cells to formation of cell aggregates containing a combination of poorly-interconnected rounded cells and spindle-shaped cells with filopodia (shCon panels of Fig. 2C) and increased cell-to-substrate adhesiveness (shCon bar of Fig. 2D) in treated control cells. Silencing of CypB almost completely prevented TGFβ-induced morphological changes (Fig. 2C) and strongly hampered the increase cell-to-substrate adhesion induced by TGFβ treatment, while CypA silencing had no effect (Fig. 2D).

**Figure 2.**
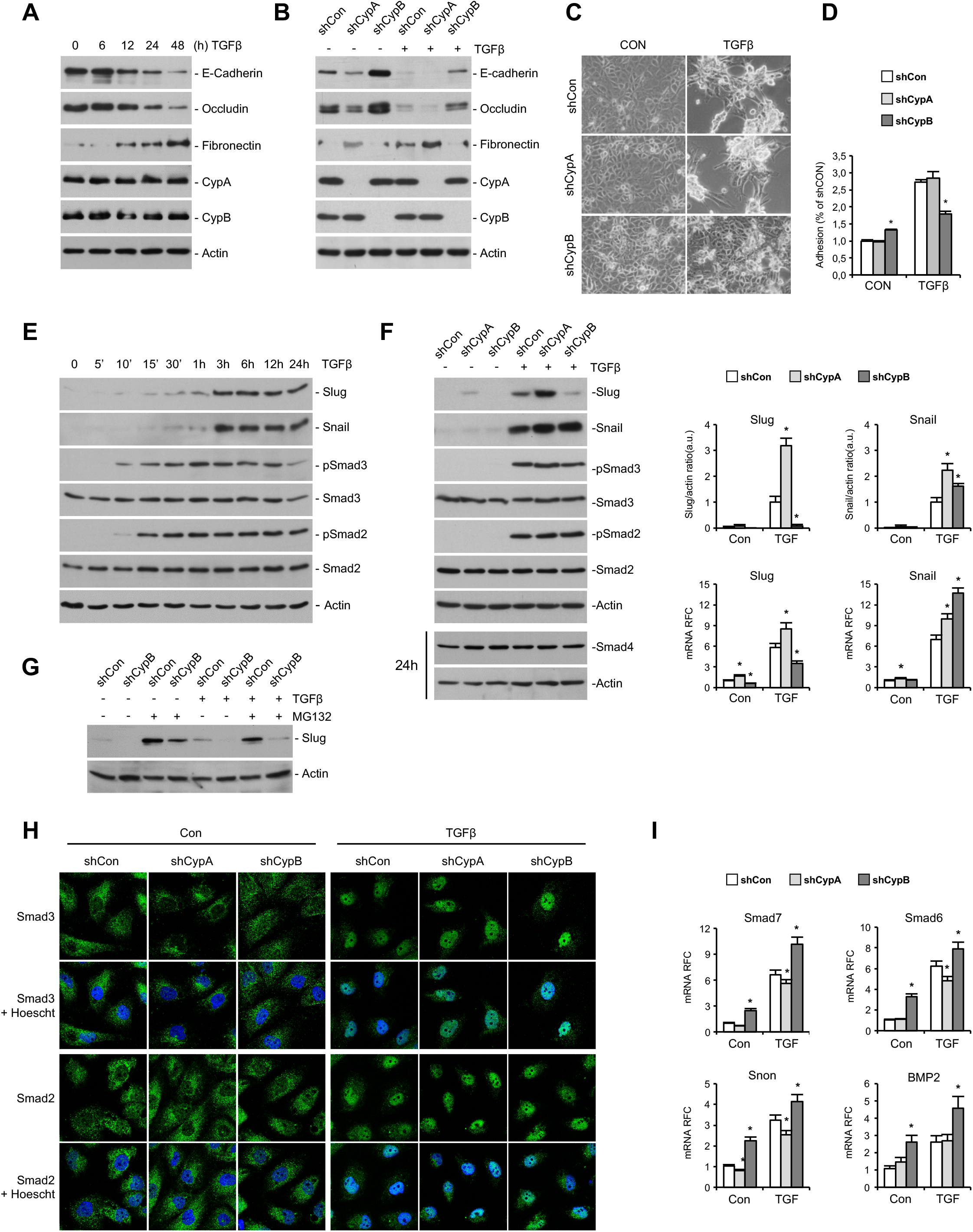
Cyclophilin A and B exert divergent effects on TGFβ action on epithelial cells. (A) HK-2 wt cells were treated with 1,5 ng/ml TGFβ for the indicated times and the expression levels of E-cadherin, occludin, fibronectin, actin, CypA and CypB were analyzed by western blot. (B) CypA- and CypB-silenced HK-2 cells were treated with 1,5 ng/ml TGFβ for 24 h and the expression levels of E-cadherin, occludin, fibronectin, actin, CypA and CypB were analyzed by western blot. (C) CypA- and CypB-silenced HK-2 cells were treated with 1,5 ng/ml TGFβ for 24 h and cells were then visualized under a light microscope. (D) CypA- and CypB-silenced HK-2 cells were treated with 1,5 ng/ml TGFβ for 48 h, trypsinized and seeded again. Cell were allowed to adhere for 30 minutes, after which unattached cells were removed and the amount of the remaining attached cells measured as indicated in methods. (E) HK-2 wt cells were treated with 1,5 ng/ml TGFβ for the indicated times and the expression levels of Slug and Snail and the phosphorylation status and expression levels of Smad3 and smad2 were analyzed by western blot. (F) Control, CypA and CypB-silenced HK-2 cells were treated with 1,5 ng/ml TGFβ for 3h or 24 h and the expression levels of slug and snail, and the phosphorylation status and expression levels of Smad3, smad2 and smad4 were analyzed by western blot. Panels on the right show the Slug/actin and Snail/actin ratios, referred to control shRNA-treated cells exposed to TGFβ, and the mRNA levels of Slug and Snail analyzed by qPCR. (G) CypA and CypB-silenced HK-2 cells were treated with 1,5 ng/ml TGFβ for 24 h, with the proteasome inhibitor MG132 (5 μM) added to cells for the lasts 16 h of TGFβ treatment. Slug levels were analyzed by Western blot. (H) Nuclear translocation of Smad3 and Smad2 in CypA and CypB –silenced cells after treatment or not with 1,5 ng/ml TGFβ for 3 h was detected by immunofluorescence using antibodies against total Smad3 and Smad2 (green).The nuclei were stained with Hoechst (blue). (I) The expression levels of Smad7, Smad6, Snon and BMP-2 in CypA- and CypB-silenced HK-2 cells treated with 1,5 ng/ml TGFβ for 24 h were analyzed by qPCR using specific proves indicated in supplemental materials. Student’s t-test was used to compare shCon vs shCypA or shCon vs shCypB for control or TGFβ treated cells. * P<0.05.

TGFβ signaling to Slug and Snail is canonically transduced by Smad proteins. Accordingly, we analyzed whether the distinct phenotypical changes observed in TGFβ-treated CypA and CypB-silenced cells could be related to Smad-dependent regulation of Slug and Snail levels. Treatment of HK-2 cells with TGFβ induced a time-dependent expression of Slug and Snail that was preceded by Smad2/3 activation (Fig. 2E). Silencing of CypB prevented TGFβ-induced slug expression and enhanced Snail expression, while silencing of CypA increased both Slug and Snail levels (Fig. 2F left). Again, these results support a predominant effect of Slug downregulation over Snail upregulation in the TGFβ-induced morphological changes observed in CypB-silenced cells. Interestingly, cyclophilin modulation of Slug and Snail occurred without changes in TGFβ-induced phosphorylation of Smad2/3 or in the expression levels of Smad4. Although to a lesser extent, these changes were also observed at the transcriptional level (Fig. 2F right). Slug and Snail proteins have a rapid turnover that is regulated by ubiquitin-mediated proteasomal degradation. To further demonstrate that CypB silencing diminished Slug expression primarily through transcriptional effects, cells were pre-treated with the proteasome inhibitor MG132 before treatment with TGFβ. As depicted in Figure 2G, treatment with MG132 alone increased Slug levels well over those of TGFβ induction. This increase was clearly reduced in CypB silenced cells, an effect even more evident after TGFβ induction, supporting the concept that CypB modulates Slug at the mRNA level.

In addition to phosphorylation, TGFβ-induced Smad2/3 signaling is also regulated by other mechanisms such as nuclear translocation or through the action of inhibitory Smad7 and Smad6, which negatively regulate TGFβ signaling by establishing an autoregulatory negative feedback loop (33). First we analyzed Smad3 and Smad2 translocation to the nucleus in CypB and CypA-silenced cells after TGFβ treatment (Figure 2H). Our results showed that neither CypB nor CypA silencing affected TGFβ-induced Smad2/3 translocation. We next analyzed the expression levels of Smad7 and 6, and of Snon, a transcriptional repressor of Smad2/3 regulated genes. As shown in Figure 2I, Smad7, Smad6, and Snon levels were transcriptionally induced by TGFβ treatment. Interestingly, all three genes were upregulated in CypB-silenced cells in both untreated and TGFβ treated cells and slightly reduced in CypA-silenced cells after TGFβ treatment.

In order to explore the mechanisms by which CypB regulates inhibitory Smads, we analyzed the levels of BMPs 2 and 7, which counteract TGFβ-induced EMT in a Smad dependent way. We were unable to detect BMP7 mRNA in HK2 cells despite the use of multiple different probes. By contrast, we detected BMP2, which was upregulated in CypB silenced cells (Fig. 2I).

### Slug modulation by CypB is independent of the CD147 receptor and extracellular CypB

Proximal tubular cells (PTC) actively contribute to the production of inflammatory mediators (5, 34, 35), thereby participating to dynamic interplay between fibrosis and inflammation. It has also been reported that NFκB, which plays a key role in inflammation, upregulates Snail both at the transcriptional level and stabilizing Snail protein (36). We hypothesized a putative role of cyclophilins on these processes and analyzed the promoter activity of NFKB in CypA and CypB silenced cells in basal conditions and upon activation by cotransfection with p65 subunit. Our results show no significant differences in NFκB promoter activity between control and silenced cells in basal conditions (Fig. 3A). However, when NFκB was activated by p65 cotransfection, we observed that the promoter activity was strongly increased in CypA-silenced cells and considerably decreased in CypB-silenced cells.

**Figure 3.**
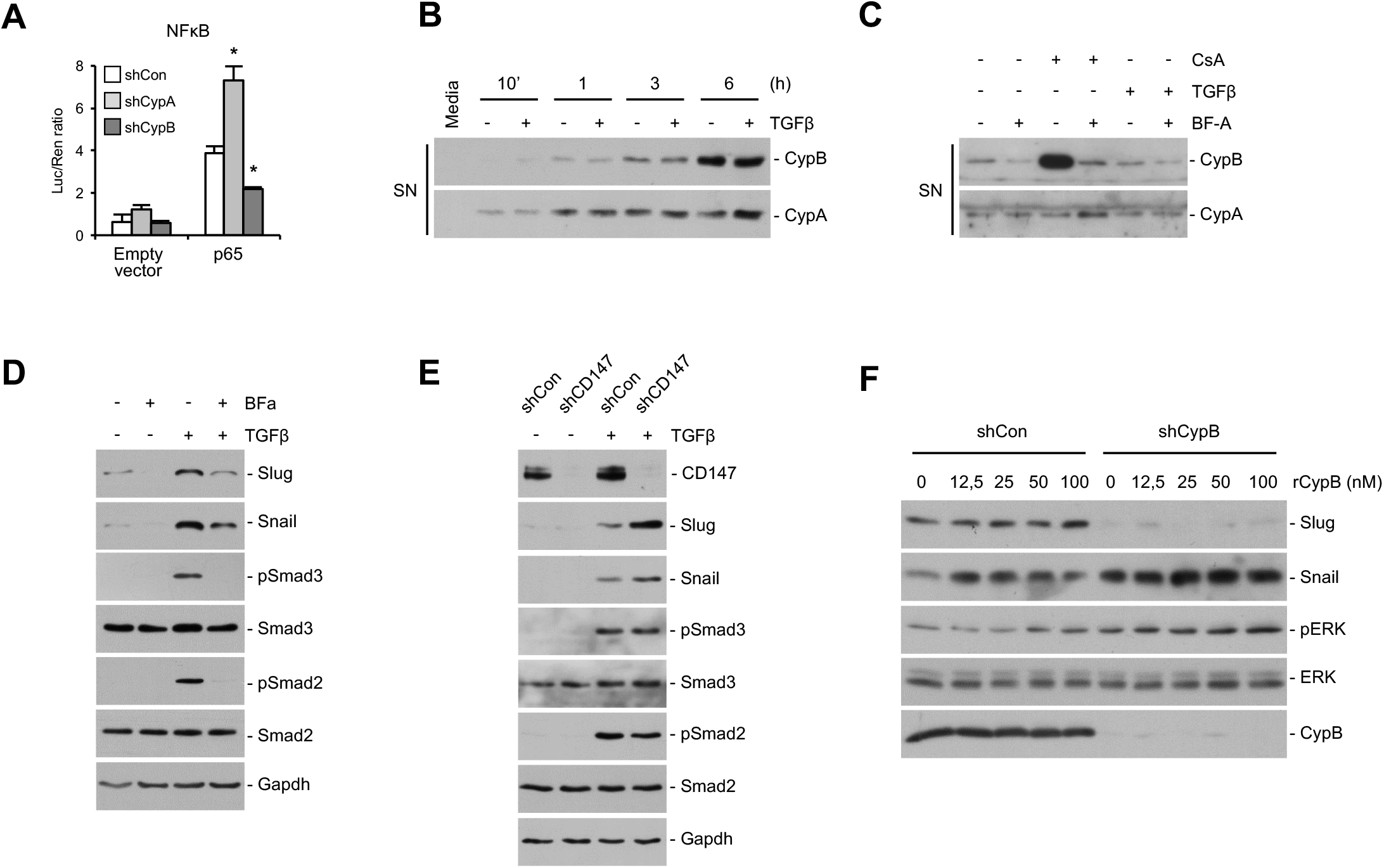
Slug modulation by CypB is independent of the CD147 receptor and extracellular CypB. Since extracellular CypA and CypB have been proposed as inflammatory mediators, we explored a potential autocrine loop in Cyp-mediated regulation of Slug and Snail. (A) NFκB activity was analyzed by cotransfecting cyclophilin-silenced HK-2 cells with plasmids containing the firefly luciferase gene under the NFκB promoter and the renilla luciferase gene under the thymidine kinase promoter, plus a negative control empty plasmid pCMV-HA (empty) or the p65 subunit of NFκB (p65). NFκB activity was analyzed using a luciferase assay kit. Transfection efficiency was normalized by the value of cotransfected Renilla luciferase. (B) HK-2 cells were treated with 1,5 ng/ml TGFβ for the indicated times, and the presence of CypA and CypB in the extracellular medium was analyzed by western blot. Culture medium that has not been in contact with cells was used as a negative control. (C) HK-2 cells were treated with 1 µM Brefeldin-A (Bf-A) for 30 minutes before addition of either 1,5 ng/ml TGFβ or 0,5 µM Cyclosporine-A (CsA) for 3 hours. Extracellular media were then collected and analyzed by western blot. (D) HK-2 cells were treated with 1 µM Brefeldin-A (Bf-A) for 30 minutes before addition of 1,5 ng/ml TGFβ and the expression levels of Slug and Snail and the phosphorylation status and expression levels of ERK1/2 analyzed by western blot. (E) HK-2 cells were stably silenced for CD147 as indicated in Methods, and treated or not with 1,5 ng/ml TGFβ for 3 hours before analyzing the levels of slug and snail and the the phosphorylation status and expression levels of Smad3 and Smad2. (F) Control and CypB-silenced HK-2 cells were treated with increasing doses of recombinant CypB for 3 h and the expression of Slug and Snail and the phosphorylation status and expression levels of ERK1/2 analyzed by western blot. Student’s t-test was used to compare shCon vs shCypA or shCon vs shCypB for control or p65 transfected cells. * P<0.05.

Because it has been described a role for extracellular cyclophilins in inflammation (37), we next aimed to explore whether cyclophilin regulation of Slug and Snail levels could be related to modulation of this inflammatory pathway in an autocrine manner. Figure 3B shows that both CypB and CypA were progressively secreted into the media of HK-2 cells and that secretion was unaffected by TGFβ treatment. To explore whether extracellular CypB could be supporting Slug expression, we blocked CypB secretion with Brefeldin-A (Bf-A) to inhibit protein transport from the endoplasmic reticulum to the Golgi apparatus as well as with Cyclosporine-A (CsA) to further induce CypB secretion. Our results show that Bf-A blocked both basal and CsA-induced CypB secretion (Fig. 3C). By contrast, neither CsA nor Bf-A had any effect on CypA secretion. This shows that CypA and CypB are secreted to the extracellular media through different mechanisms. Bf-A pre-treatment prevented basal and TGFβ-induced Slug expression (Fig. 3D) but, contrarily than CypB silencing, also reduced Snail expression and Smad3 and Smad2 activation. To further explore the involvement of extracellular cyclophilins, we knocked down CD147, which is, to our knowledge, the only known receptor for extracellular CypB and CypA. Our results show that CD147 silencing increased, rather than decreased, Slug and Snail levels, thereby resembling our findings in CypA but not in CypB-silenced cells (Fig. 3E). Finally, and to further investigate whether extracellular CypB could be modulating Slug levels in a CD147 independent manner, CypB-silenced cells were treated with increasing doses of recombinant CypB. Our results show that exogenous CypB was unable to restore Slug expression or downregulate Snail levels to levels of non-silenced cells (Fig. 3F). ERK1/2 phosphorylation was used as a control of CypB treatment effectiveness (38). Taken together these results argue against and autocrine loop in Slug modulation by CypB.

### Slug regulation by CypA and CypB depends on PPIase activity of CypA and ER location of CypB

To further investigate the mechanisms underlying modulation of Slug and Snail expression by CypB and CypA, we re-introduced mutated forms of CypB and CypA lacking PPIase activity or, in the case of CypB, its N-terminal signal peptide, in the corresponding silenced-cells (Fig. 4A). To do so, we first generated a shRNA-resistant wild-type CypB (wt CypB) by introducing silent mutations into the shRNA-targeted sequence for CypB to allow its escape from degradation by the RISC complex. ShRNA against CypA was directed to the 3’UTR, and thus reintroduction of a wild-type form of CypA (wt CypA) did not require any further modification. Re-introduction of the wt forms served as rescue experiments to validate that the effects observed in silenced cells were not due to off-targets effects. Over these constructs, we mutated critical residues for PPIase activity (ΔPPI mutants) in both CypB (R62A) and CypA (R55A), and separately deleted the signal peptide (Δ(K2_A25)) directing CypB to the endoplasmic reticulum (ΔER mutant). Finally, an HA-tag was added at the C-terminus of each cDNA. All these constructs were stably transduced into cyclophilin-silenced cells using lentiviral particles and selected with a different antibiotic than the one used for silencing selection. Western blot assays demonstrated that the wild-type and the mutant forms of both CypA and CypB were successfully reintroduced in the corresponding silenced cells (Fig. 4B and 4E, respectively). Moreover, by using confocal IF we observed that both wt and ΔPPI forms of CypB staining correlated with ER location, as determined by the ER transmembrane protein calnexin (CNX), and were successfully secreted to the extracellular medium (Fig. 4F and 4E, SN panel), while CypB ΔER did not (Fig. 4F and 4E, SN panel).

**Figure 4.**
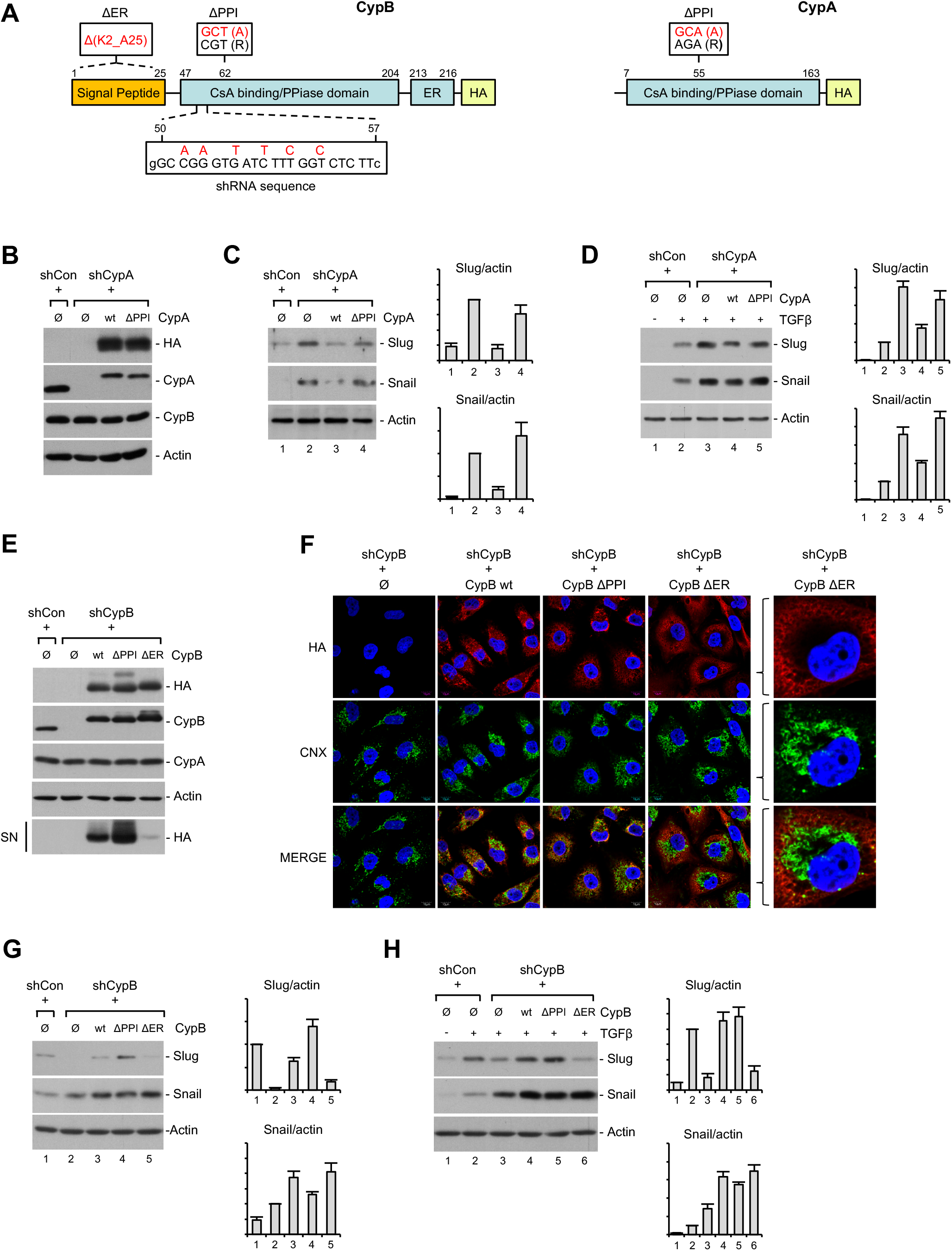
Slug regulation by CypA and CypB depends on PPIase activity of CypA and ER location of CypB. shRNA-resistant wild-type (wt) or shRNA-resistant mutants defective in PPIase activity (ΔPPI) or, in the case of CypB, its signal peptide (ΔER), were cloned into lentiviral expression vectors and reintroduced into CypA or CypB-silenced cells to discard off-targets effects of shRNA and to study the involvement of PPIase activity and ER location. (A) Schematic diagram summarizing the different mutations introduced in CypA and CypB expression vectors. (B) The expression levels of CypA wt and mutant forms reintroduced into CypA-silenced cells were determined by Western Blot. Ø corresponds to the empty expression vector. (C and D) Western Blot analysis showing that the increase in Slug and Snail levels observed in CypA-silenced cells is prevented by reintroducing HK-2 CypA wt but not the R55A mutant, either in untreated cells (C) or cells treated with 1,5 ng/ml TGFβ for 3 hours (D). (E) The expression levels of CypB wt and mutant forms reintroduced into CypB-silenced cells were also analyzed Western Blot in cell extracts and cell supernatants (SN). Ø corresponds to the empty expression vector. (F) Immunofluorescence staining of reintroduced CypB wt and the ΔPPI and ΔER mutants of CypB. CypB wt and ΔPPI colocalized with the ER marker calnexin (CNX) while CypB ΔER did not. (G and H) Western Blot analysis of the expression levels of Slug and Snail, after reintroduction of CypB wt or ΔPPI and ΔER mutants, either in untreated cells (G) or in cells treated with 1,5 ng/ml TGFβ for 3 hours. Actin ratios are referred to control shRNA-treated cells for Slug and Snail.

As shown in Figure 4C, reintroduction of wt CypA into CypA-silenced cells decreased Slug and Snail levels to those of control cells, while reintroduction of the ΔPPI mutant of CypA failed to do so, indicating that the PPIase activity of CypA is necessary to maintain Slug and Snail at basal levels. These results were also observed in the presence of TGFβ (Fig. 4D). On the other hand, reintroduction of CypB-wt into CypB-silenced cells increased Slug levels almost to control levels while that of the ΔPPI mutant not only rescued but increased Slug levels over those of control cells (Fig. 4G). Finally, reintroduction of the ΔER mutant failed to restore Slug levels. Again, these effects were observed in the presence of TGFβ (Fig. 4H). These results reinforce the idea that CypB is required for Slug expression and that its presence in the ER but not its PPIase activity is required for this effect.

### CypB modulates Slug levels through calcium regulating elements in the ER

As shown above, ER location of CypB is mandatory for Slug expression. Since it has been previously described that CypB interacts with the calcium-related ER chaperones calreticulin (CRT) and calnexin (CNX) (39), and CRT has been involved in the regulation of Slug expression (40–42), we analyzed whether the modulatory effects of CypB on Slug levels could be related to changes in CRT/CNX localization or altered interaction with these chaperones. Figure 5A shows that CRT and CNX colocalized discretely, fitting with their ER luminal and transmembrane location, respectively. We also observed that localization of CRT and CNX was unaffected by either CypA or CypB silencing. Next, wild type, PPIase and signal peptide defective mutants of CypB were re-introduced in CypB-silenced cells and immunoprecipitated with HA antibody. As shown in Figure 5B, CRT immunoprecipitated with CypB wt and to a higher extent with CypB ΔPPI but no interaction was detected with the CypB ΔER mutant. Interestingly, this interaction pattern paralleled that of Slug expression shown in Figure 4G. By contrast, we detected CNX interaction with the CypB ΔER mutant but not with the wt form. Although a strong interaction of CNX with CypB ΔPPI was also observed, it is worth mentioning that CNX levels were increased in CypB ΔPPI cell extracts (input). Taken together these results suggest that a CypB-CRT complex rather than a CypB-CNX complex could be mandatory for CypB regulation of Slug. They also indicate that interaction with either CRT or CNX depends on CypB location.

**Figure 5.**
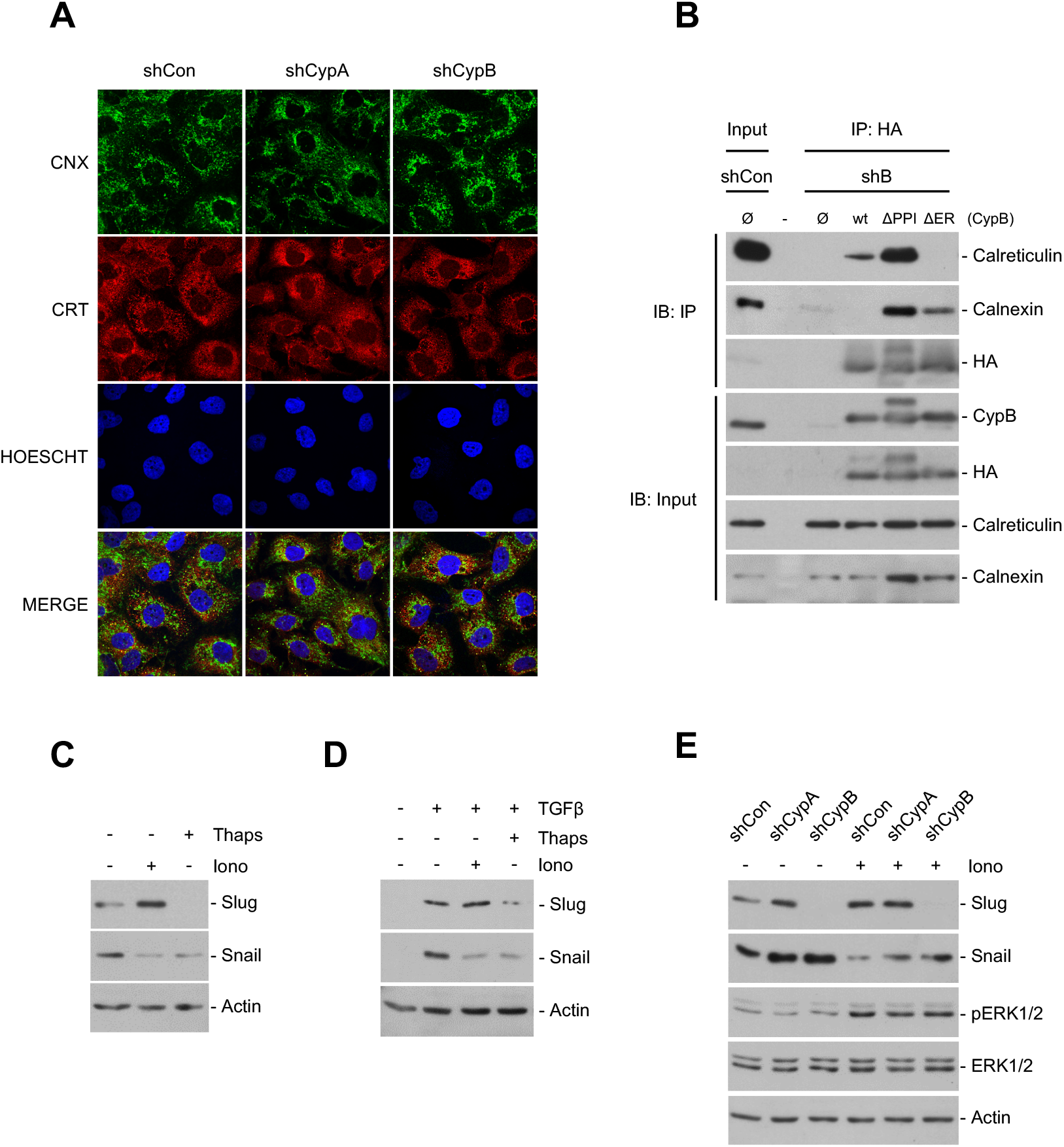
CypB could be modulating Slug levels through calcium regulating elements in the ER. To explore whether the modulatory effects of CypB on Slug levels could be related to calreticulin (CRT) and calnexin (CNX), we analyzed CRT and CNX subcelular localization and protein interactions in HK-2 silenced cells. (A) Immunofluorescence staining of CRT (ER luminal) and CNX (ER transmembrane) was unaffected by either CypA or CypB silencing. (B) Wild type, PPIase and signal peptide defective mutants of CypB were re-introduced in CypB-silenced cells and immunoprecipitated with HA antibody, followed by immunoblotting with CRT, CNX, HA or CypB. (C) To explore if the changes in slug levels could be related to calcium homeostasis disturbance, HK-2 cells were treated with 5 µM thapsigargin or 1 µM Ionomycin for 3 hours and Slug and Snail levels were analyzed by western blot. (D) HK-2 cells were treated with 5 µM thapsigargin or 1 µM Ionomycin for 30 minutes before addition of 1,5 ng/ml TGFβ for 3 hours. (E) Control, CypA and CypB-silenced cells were treated with 1 µM Ionomycin for 3 hours and Slug and Snail levels and the phosphorylation status and expression levels of ERK1/2 analyzed by western blot.

CRT is an important ER calcium buffering protein involved in regulating ER calcium storage and release. To ascertain if Slug downregulation in CypB silenced cells could be due to ER-related calcium signaling, we analyzed whether Slug and Snail levels were modulated by alterations of ER calcium stores. ER calcium pools were depleted by exposing cells to thapsigargin or ionomycin (Fig. 5C). Our results show that basal Slug levels were increased by ionomycin but suppressed by thapsigargin, while Snail levels were downregulated by both. These effects were also observed in the presence of TGFβ (Fig. 5D). Interestingly, ionomycin was unable to induce Slug expression in CypB-silenced cells, and the ionomycin-induced decrease of Snail levels was only partially attenuated in both CypB and CypA-silenced cells. These changes occurred independently of ionomycin-induced ERK activation. Taken together these results suggest that CypB could be regulating Slug levels through ER calcium-related events.

### CypB depletion ameliorates inflammation and fibrosis after UUO

To expand on the results performed in vitro and to assess the potential contribution of CypB in the development of fibrosis, global CypB-KO mice and wild-type (wt) littermates (Fig. 6A) were subjected to unilateral ureteral obstruction (UUO) of the left kidney for one-week period to mimic the pathological conditions underlying early events in renal fibrosis (Fig. 6B). Controls corresponded to contralateral (non-ligated) right kidneys from the same mice. Our results show that while no apparent differences were observed between the non-ligated kidneys from wt and CypB KO mice regarding overall kidney morphology (HE), inflammation (F4/80) and collagen deposition (MT), we found that obstructed kidneys from CypB KO mice were protected from the tubular distension, inflammation and fibrosis observed in wt mice under UUO (Fig. 6C). Sham-operated mice were also included and results were the same as those found in non-ligated kidneys (not shown). These results suggest that CypB deficiency is associated to reduced UUO-induced kidney fibrosis.

**Figure 6.**
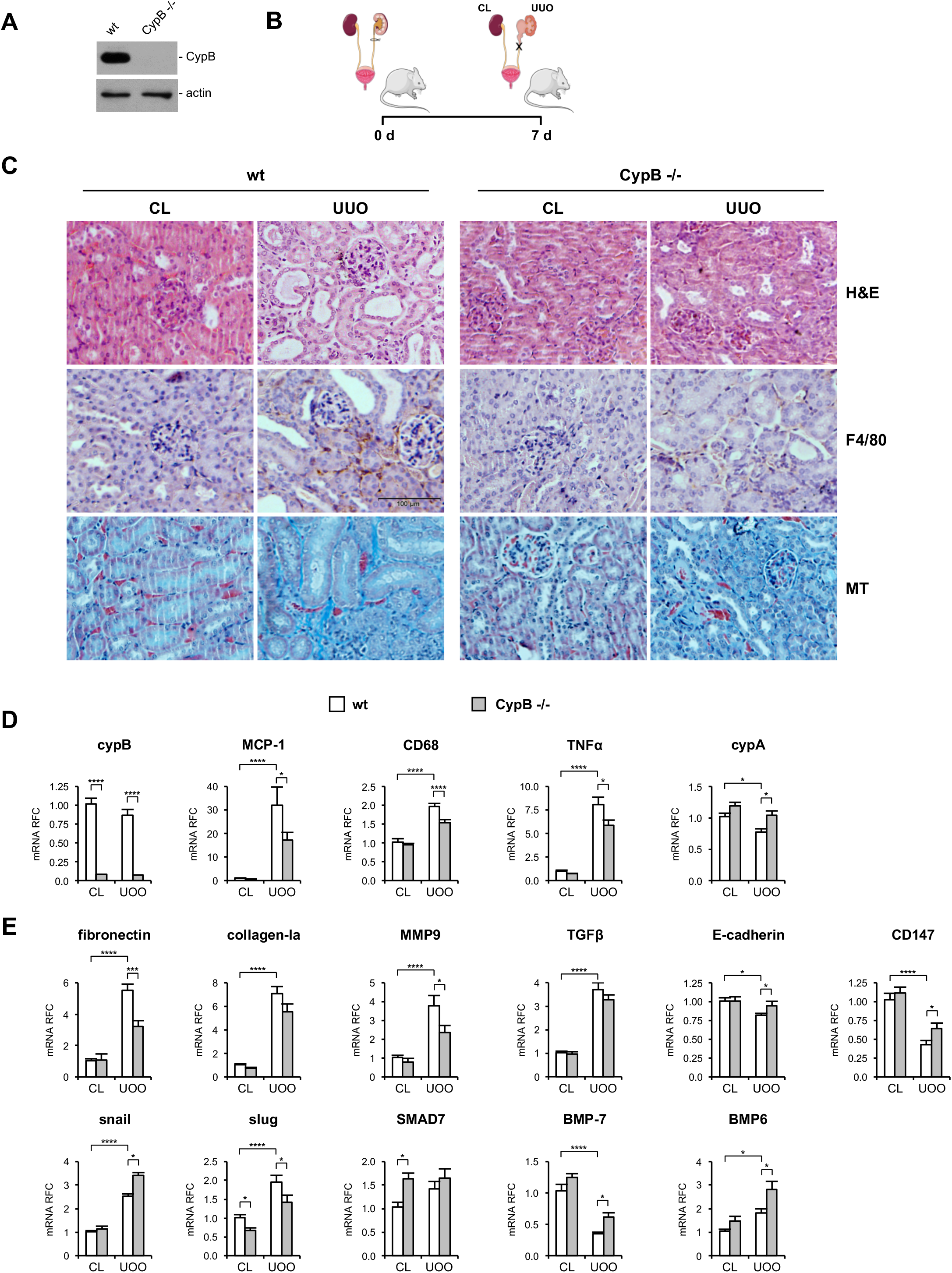
CypB depletion ameliorates inflammation and fibrosis after UUO. CypB KO mice were used to assess the involvement of CypB in kidney fibrosis. (A) The expression levels of CypB were analyzed by Western Blot in kidneys from CypB KO (CypB -/-) mice and control wild type littermates (wt). (B) Scheme depicting the experimental approach: CypB-KO mice and control littermates were subjected to unilateral ureteral obstruction as a model of renal fibrosis. Mice were sacrificed 7 days after obstruction and total RNA was extracted from the right contralateral kidneys (CL) and left obstructed kidneys (UUO). (C) Hematoxylin and eosin (H&E), mouse macrophage marker F4/80, and Masson’s trichrome (MT) staining of kidneys sections from contralateral kidneys (CL) and obstructed kidneys (UUO) of wt and CypB KO mice. (D) CypB, MCP1, CD68, TNFα and CypA mRNA levels detected by qRT-PCR. (E) Fibronectin, collagen-Ia, MMP9, TGFβ, E-cadherin, CD147, snail, slug, Smad7, BMP-7, and BMP-6 levels detected by qRT-PCR. Each column shows mean ± SEM; n = 8 animals per group. Data were analyzed by two-way analysis of variance (ANOVA) followed by Tukey multiple comparison test. ns P ≥ 0.05; * P<0.05, ** P<0.01, *** P<0.001; ****P<0.0001.

Since UUO induces a robust inflammatory response we compared the levels of pro-inflammatory cytokines present in obstructed kidneys of wt and CypB KO mice (Fig. 6D). We found that in wt mice the levels of tumor necrosis factor α (TNF-α), macrophage chemo-attracting protein 1 (MCP-1) and the pan-macrophage marker CD68 were strongly up-regulated in the obstructed kidneys in comparison to the right contralateral kidneys. This increase was significantly reduced in the kidneys of CypB KO mice. In non-obstructed kidneys, there were no differences in the expression of these inflammatory markers between wt and CypB KO mice. In accordance with the protein levels shown in Figure 6A, CypB mRNA levels were almost undetectable in CypB KO mice and were not affected by the UUO (Figure 6D). We also observed that the genetic deletion of CypB prevented the down-regulation of CypA induced by UUO.

To further investigate the effects of CypB deletion in kidney fibrosis we analyzed the levels of fibrosis and EMT related genes. In obstructed kidneys of wt mice, we observed a potent and significant increase in the expression levels of ECM components fibronectin and collagen-Ia and the metalloproteinase MMP-9, as well as of TGFβ. We also observed a significant reduction of E-cadherin and CD147 mRNA levels in comparison with those found in the contralateral kidneys (Fig. 6E). Our results show that genetic deletion of CypB significantly prevented the UUO-induced increase of fibronectin and MMP-9 and the decrease of E-cadherin and CD-147 observed in wt mice. However, we didn’t observe significant differences in TGFβ levels between wt and CypB KO mice that could explain the aforementioned protective effects of CypB knockdown. To gain further insights into the mechanism of action of CypB, we next investigated the levels of the TGFβ downstream mediators Snail and Slug, and the levels of the antifibrotic factors SMAD7, BMP7 and BMP6, which counteract the TGFβ pathway. Kidney obstruction increased the levels of Slug, Snail, SMAD7 and BMP6 while reduced those of BMP-7 (Fig. 6E). CypB knockdown reduced Slug levels both in contralateral and obstructed kidneys, while increased those of Snail only after UUO. CypB KO mice also show a reduced decrease and an augmented increase in BMP7 and BMP-6, respectively, after UUO. Finally, we also observed that SMAD7 levels were upregulated in non-ligated kidneys of CypB KO mice.

These results, together with those found in HK-2 cells, indicate the association of CypB deficiency and a lower fibrotic and inflammatory response after UUO and suggest a potential role of CypB on EMT processes.

## Discussion

It is increasingly accepted that after sustained kidney injury, tubular epithelial cells undergo a partial EMT or type 2 EMT, which contributes to kidney fibrosis. This process appears to be plastic, where cells are not engaged in an unidirectional process or locked into one differentiated state, but eventually transit back to the epithelial phenotype (43). Accordingly, the identification of factors regulating this epithelial plasticity could improve the understanding of kidney fibrosis. In the present work, we demonstrate for the first time that CypB is a critical regulator of tubule epithelial cell plasticity on the basis that: i) CypB silencing caused epithelial differentiation in the proximal tubule-derived cell line HK-2, ii) CypB silencing prevented TGFβ-induced EMT in HK-2 cells, and iii) Global CypB knockdown is associated to a reduced UUO-induced kidney fibrosis. Interestingly, silencing of CypA in HK-2 cells exerted almost diametrically opposite effects on epithelial cells than CypB silencing. All these effects most likely result from CypB and CypA regulation of the transcriptional repressor Slug and Snail.

It is becoming increasingly evident that the role of cyclophilins is not restricted to protein folding and that they are also involved in multiple cellular processes such as cell division, protein trafficking, cell signaling, transcriptional regulation and stress tolerance among others (44). Results presented in this study show that in a global CypB knockdown the levels of inflammation and fibrosis mediators were reduced and that the loss of E-cadherin in mouse obstructed kidneys was prevented. Since a pro-inflammatory role of extracellular CypB has been described (45), it could not be discarded that the protective effects of CypB knockdown on kidney fibrosis were secondary to a reduction in inflammation. Our results in HK-2 cells not only support a direct effect of CypB silencing on promoting epithelial phenotype but also in reducing inflammation through NFκB inhibition. Altogether, our results suggest that the gain of epithelial markers observed in CypB silenced proximal tubule cells might resemble the mesenchymal to epithelial (MET) transition undergone by surviving tubule cells upon injury, thus promoting the functional and histological features that would return the kidney to normal function. Accordingly, it could be expected that CypB overexpression recapitulated events driving to a more de-differentiated phenotype. Actually, it has been described that CypB overexpression is related to malignant progression in several types of tumors including breast cancer, hepatocellular carcinoma, gastric tumors and malignant gliomas (46–49). Contrarily to the effects caused by CypB silencing, knockdown of CypA induced a more undifferentiated phenotype regarding the molecular and functional epithelial markers studied. These results are in agreement with previous studies showing loss of cell-cell contacts and increased fibronectin expression after CypA-silencing of human renal epithelial cells from nephrectomy specimens (50). Considering the above phenotypical consequences of CypA silencing in cultured cells, the reduction in CypA levels observed in mouse kidneys after ureteral obstruction could be relevant for the development of fibrosis. In this sense, the increased levels of CypA in injured kidneys of CypB KO mice may respond to a physiological compensatory mechanism. In summary, results obtained in both, CypB KO mice and in cultured proximal tubule cells demonstrate a role for CypA and CypB in epithelial plasticity and homeostasis; while CypB would foster a more dedifferentiated state, CypA would preserve the epithelial phenotype.

These striking opposite actions of CypB and CypA silencing on the HK-2 epithelial phenotype revealed, for the first time, a differential regulation of Snail and Slug transcriptional repressors by these cyclophilins. Specifically, CypA-silenced cells showed upregulated levels of both Snail and Slug while CypB-silencing strongly downregulated Slug and upregulated Snail; thereby indicating that while CypB represses Snail but is required for Slug expression, CypA acts as a repressor of both Slug and Snail proteins. Moreover, we can conclude that, at least in CypB silenced cells, loss of Slug drives the epithelial cell phenotype in spite of increased Snail levels. In this sense, and despite their high homology in their DNA binding and SNAG domains, Snail and Slug present common and nonequivalent functions in epithelial promoter repression, DNA binding and EMT-inducing ability (51). This predominant role of Slug is also particularly relevant since Snail has been getting most of the attention and considered as a sufficient and necessary factor to induce EMT and fibrosis in mouse kidney (11).

Snail and Slug are considered the master-regulators of TGFβ-induced EMT and kidney fibrosis. In cultured proximal tubular cells, including HK-2, treatment with TGFβ is enough to induce Slug and Snail and trigger EMT. Consistent with our results in untreated cells, CypB silencing prevented TGFβ-induced Slug expression and its profibrotic effects, while CypA silencing enhanced them. TGFβ regulates renal fibrosis positively by receptor regulated R-Smads (Smad2/3), but negatively by inhibitory I-Smads (Smad6/7) which are transcriptionally induced by TGFβ establishing an important negative feedback loop. In this sense, blockade of TGFβ signaling by Smad7 gene therapy is known to prevent experimental renal fibrosis (52). Interestingly, it is worth noticing that, from the gene panel analyzed, Smad7 and Slug were the only genes differentially expressed in non-ligated kidneys of CypB KO in comparison with wt mice; where Smad7 mRNA overexpression was associated with Slug down-regulation. In agreement with that, we found that in TGFβ treaded HK-2 cells, Smad7 and also Smad6 expression was upregulated in CypB-silenced cells. I-Smads antagonize TGF-β signaling through multiple negative feedback mechanisms that include: i) formation of Smad7 stable complexes with activated type I TGFβ receptor ALK5/TβRI which blocks the phosphorylation of R-Smads and subsequent nuclear translocation of R-Smad/Smad4 heterocomplexes; ii) by recruiting ubiquitin E3 ligases, such as Smurf1/2, resulting in the ubiquitination and degradation of TβRI and iii) in the nucleus, by interfering with the formation of the functional R-Smad/Smad4-DNA complex on target gene promoters (53). On this basis and considering that CypA and CypB silencing modulate Slug and Snail levels upon TGFβ treatment without changes in either Smad2/3 phosphorylation or nuclear translocation, our results would suggest a direct inhibitory effect of Smad7 over Slug promoter in CypB-silenced cells. This mechanism seems to be specific for Slug promoter, since it is not affecting Snail expression. Besides TGFβ, Smad7 expression is also regulated by BMPs. BMPs further counteract TGFβ pathway through other mechanisms such as control of Snon expression, which represses TGFβ signaling by inactivating Smad transcriptional complexes. Considering that levels of BMP7 and 6 in CypB KO mice and BMP-2 and Snon in CypB-silenced HK-2 cells were upregulated, and that CypB knockdown had no significant effect on UUO-induced TGFβ levels, our results suggest that lack of CypB would impact Slug expression by counteracting, rather than hampering, TGFβ signaling.

Mechanistically, both CypA and CypB seem to be acting on Slug and Snail through different mechanisms. Thus, while the PPIase activity of CypA is required to keep both Slug and Snail downregulated, the presence of CypB signal peptide, but not its PPIase activity, is mandatory to allow slug expression. These results fit with a previous report showing that CypA, through its PPIase activity, participates in epithelial differentiation of kidney intercalated cells by mediating matrix assembly of the extracellular matrix protein hensin (54).The involvement of the PPIase activity of CypA in the maintenance of epithelial phenotype was also supported by the fact that, although cyclosporine-A (CsA) inhibits the PPIase activity of both CypB and CypA, silencing of CypA, rather than that of CypB, mimics the CsA-induced phenotypic changes previously described on proximal tubular cells (13), pointing to CypA as the main target of CsA effects. Regarding CypB, the involvement of its signal peptide on slug expression might either require progression through the secretory pathway ultimately leading to secretion, or the need for CypB localization within the ER. Considering the aforementioned extracellular role of cyclophilins, the existence of a cyclophilin autocrine signaling loop affecting slug expression could not be discarded. Nevertheless, our results argue against an extracellular role of CypB in modulating slug since: i) no differences in either CypB or CypA secretion were observed after TGFβ treatment; ii) silencing of CD147, the only known receptor for extracellular cyclophilins, did not prevent but rather increase slug expression; and iii) exogenously added recombinant CypB was unable to restore slug levels. It is worth mentioning that the effects of CD147 silencing on Slug expression resembled those of CypA silencing, suggesting that, at least for CypA, an extracellular loop could be involved in CypA regulation of Slug and Snail. In accordance with this, it has been previously described that CypA acts as a survival-enhancing autocrine factor in mouse ESC cultures (55). Moreover, it has been reported that extracellular CypA activates the SMAD pathway and promotes inflammation in biliary atresia and that targeting this extracellular CypA ameliorates disease progression (56).

From the aforementioned results we conclude that CypB would be mediating Slug expression by means of its non-catalytic chaperone role within the ER. From the initial discovery of CypB interaction with CAML (calcium-signal modulated cyclophilin ligand) (57), novel CypB partners have been identified, including the sodium-potassium ATPase (58) and the human TRPV6 calcium channel protein (59) among others. In addition, CypB participates in macromolecular chaperones complexes in the ER lumen together with the ERp72, CRT, CNX, BiP (GRP78) and GRP94 (60, 61), and the lectin chaperones calnexin (CNX) and calreticulin (CRT) (39). The latest are of special interest because their involvement in calcium signaling and to previous reports indicating that CRT was required for TGFβ-induced EMT (41, 42, 62–64). The biological and functional relevance of CypB interaction with CRT and CNX has not been still elucidated. Our results show that in HK-2 cells, CypB preferentially interacts with CRT since no interaction with CNX was observed under control conditions. Interestingly, CypB-CRT interaction was increased when the PPIase activity of CypB was muted, and was totally prevented when CypB was not present in the ER, closely paralleling that observed in Slug expression and therefore suggesting a potential role of CypB-CRT interaction in Slug regulation. According to the literature, and as we observed in CypB-silenced cells, CRT seem to be preferentially acting over Slug, since CRT over expression in kidney epithelial MDCK cells (40) or HK-2 cells (42), resulted in Slug upregulation without apparent (40), or much more reduced (42), changes in Snail levels. Moreover, as occurred in CypB silenced HK-2 cells, CRT depletion does not impact canonical TGFβ signaling as TGFβ was still able to stimulate Smad activity in CRT -/- MEFs (63). On the other hand, interaction of CypB with CNX was only detected when CypB has its PPIase activity mutated or when CypB was unable to enter the ER. CypB interacts with CRT and CNX through their P-domain, and complex formation is not affected by CsA, confirming the functional independence of the P-domain binding and PPIase activity (65). We hypothesize that, since CypB interaction with other partners might require intact PPIase residues, CRT/CNX interaction with ΔPPI CypB would benefit from a larger pool of unbound CypB. Nevertheless, interaction of CNX with ΔER CypB remains to be explained considering that CNX is an integral ER transmembrane protein and that the p-domain containing CypB interaction sites faces the lumen of the ER. Considering our results and previous data on the literature, it is entirely plausible that CypB could be modulating Slug expression through its interaction with CRT.

It has been described that CRT mediates TGFβ-dependent transcriptional responses through its role as a calcium signaling regulator, rather than through its chaperone/UPR function (64). In this sense, cells lacking CRT have impaired calcium release from the ER (64), suggesting that calcium release could be a key factor in Slug modulation. However, our results suggest that an increase in cytosolic calcium levels *per se* is not determinant in slug modulation, since treatment with ionomycin or thapsigargin (Tg), which both ultimately lead to increased cytosolic calcium, exerted opposite effects on Slug levels, by either increasing or decreasing them, respectively. Thus, it is likely that a more specific calcium event within the ER underlies Slug modulation by CypB. Similar results were obtained by Zimmerman et al. (63), showing that TGFβ treatment in the presence of ionomycin, despite an increase in cytoplasmic calcium, was still unable to increase ECM transcript in CRT-deficient cells. Results from these authors suggested that, although CRT-mediated calcium regulation is a critical factor for TGFβ-induced ECM stimulation, calcium itself is not sufficient, and other CRT-dependent factors should be involved. In a similar manner, we observed that CypB silencing prevented the increase in Slug levels after Ionomycin treatment. Ionomycin is a Ca^2+^ ionophore, while thapsigargin irreversible inhibits the SERCA pump, abrogating the reuptake of cytosolic calcium into the ER, and thus causing depletion of Ca^2+^ within the ER. Since both thapsigargin-induced SERCA inhibition and CypB silencing had the same effects on Slug expression, and it has been described that CRT regulates SERCA activity (66), we hypothesize that the lack of Slug expression on CypB-silenced cells could result from impaired CRT modulation of SERCA activity. In addition, it has been described that CypB overexpression protects against thapsigargin-induced cell death, attenuating calcium release from the ER (67). Taken together, our results indicate that CypB regulation of Slug could be related to the indirect action of CypB on ER calcium stores through interaction with the major calcium binding chaperone CRT.

In conclusion, we uncovered new roles for CypA and CypB in modulating proximal tubular cell phenotype in differentiation and EMT processes, most likely through regulation of the expression of the transcriptional repressors Slug and Snail. Our results also reconsider the functional relevance between both repressors, placing Slug as a key regulator of EMT in the fibrotic kidney. As modulators of EMT, targeting CypA or CypB could have a great impact not only on the overall outcome of kidney fibrosis but also in other processes where cell-cell contacts are critical, such as in cancer. Moreover, we also propose a crucial role of CypB in regulating the inflammatory response that precedes kidney fibrosis, establishing CypB as an important link between inflammation and fibrogenesis. Finally, development of drugs specifically targeting CypB could have a therapeutic benefit to reduce TGFβ effects and ameliorate kidney fibrosis.

## Methods

### Unilateral ureteral obstruction (UUO) procedure

Unilateral ureteral obstruction was performed as previously described (68). Briefly, mice were anesthetized by intraperitoneal injection of pentobarbital (50 mg/kg). Through a 2 cm midline abdominal incision, the left kidney was exposed by retraction of the intestines using a self-retaining microdissection retractor and the left ureter was carefully dissected from surrounding tissue. The ureter was then doubly ligated at the midpoint between the kidney and the bladder using sterile 6-0 silk suture. The surgical incision was then closed and the mouse was allowed to recover from anesthesia; postoperative analgesia (buprenorphine, 0.1 mg/kg, SQ) was administered. Sham-operated control mice underwent a similar surgical procedure without ligation of the ureter.

### RNA extraction and RT-PCR

Total RNA was isolated using TRIzol® Reagent (#15596-026, Ambion, Life Technologies) according to manufacturer’s instructions. Reverse transcription was performed from 2 μg of total RNA with kit High-Capacity RNA-to-cDNA™ Kit (Applied Biosystems, #4387406). Quantitative RT-PCR was carried out on an ABI PRISM 7900 Sequence Detection System (Applied Biosystems) using pre-designed FAM-labeled TaqMan probes (Applied Biosystems). All probes used in this work are listed in Supplemental Material. Analysis of relative gene expression data was performed using the 2^-(ΔΔCt)^ method after normalizing to TBP.

### Cell culture

Human kidney proximal tubule cells (HK-2), which retain morphologic and functional attributes of normal adult human proximal tubular epithelium (69), were cultured in medium A (DMEM:Ham’s F12 (1:1, v/v), 20 mM HEPES, 2mM L-glutamine, 12.5 mM D-glucose, 60 nM sodium selenite, 5 μg/ml transferrin, 50 nM dexamethasone, 100 U/ml penicillin and 100 μg/ml streptomycin) supplemented with 2% fetal bovine serum (FBS), 5 μg/ml insulin, 10 ng/ml epidermal growth factor (EGF) and 1 nM triiodothyronine, at 37 °C in a 95:5 air:CO_2_ water-saturated atmosphere. For all experiments, cells were seeded at 0.1×10^6^ cells/ml. For TGFβ treatments, cells were starved in medium A supplemented with only 0.1% FBS and without insulin, EGF or triiodothyronine (starvation medium). After 16 hours starvation, medium was removed and cells were treated for the indicated times with fresh starvation medium containing 1.5 ng/ml TGFβ (#100-B-001, R&D Systems). When indicated, cells were treated with the indicated doses of human TGFβ (#100-B-001, R&D Systems, Minneapolis, MN, USA), MG-132 (#BML-PI102, Enzo Life Sciences, Farmingdale, NY, USA), Cyclosporine-A (#239835, Calbiochem, San Diego, CA, USA), Brefeldin-A (#B6542, Sigma-Aldrich, St. Louis, MO, USA), human CypB (#NBC1-18424, Novus Biologicals, Littleton, CO, USA), Ionomycin (#I0634, Sigma-Aldrich) Thapsigargin (#586005, Calbiochem).

### Gene silencing

CypA, CypB and CD147 silencing was performed as described in (58). For CypA and CypB silencing, shRNA-containing lentiviral particles were generated by co-transfecting HEK293T cells with second generation vectors including the transfer vector pAPM, carrying the shRNA and puromycin resistance, the HIV-1 packaging plasmid psPAX2, and a VSVg expression plasmid (pMD2.G) (complete information about these vectors is available at (70)). For CD147 silencing, a MISSION® TRC shRNA transfer vector (TRCN0000006732) was cotransfected with the third generation vectors VSVG, RTR2 and PKGPIR, which provide the envelope, packaging and reverse-expressing proteins, respectively. Viral supernatants were then harvested, supplemented and added to HK-2 cells in the presence of polybrene (Hexadimethrin bromide, Sigma-Aldrich). shRNA sequences were indicated in supplemental material.

### Co-Immunoprecipitation

For coimmunoprecipitation (CoIP) assays, cells were lysed in CoIP buffer (50 mM Tris-HCl pH 7.5, 150 mM NaCl, 2 mM EDTA, 10 % glycerol and 1 % Triton) containing protease and phosphatase inhibitors, and 750 µg of cell extracts were incubated with 40 µl of anti-HA Affinity Matrix (clone 3F10, #11815016001, Roche, Mannheim, Germany) overnight at 4 °C on a rotary mixer. Beads were then washed 5x with CoIP buffer, and proteins bound to the beads were eluted by incubating the beads with 80 µl of 1 µg/µl HA peptide (#11666975001, Roche) for 1 hour at 37 °C.

### Western blot

Cells were lysed with RIPA buffer supplemented with protease inhibitor cocktail (Sigma-Aldrich) and the protein content of cellular extracts was quantified by the BCA assay (Thermo Scientific). Equal amounts of whole cell extract protein were run on SDS PAGE gels and transferred onto PVDF membranes. Membranes were then blocked 1 hour at RT and incubated overnight at 4 °C with the corresponding antibodies (a complete list of all antibodies used in this work is shown in Supplemental Material). Finally, membranes were developed with the enhanced chemiluminescence method (Millipore) and exposed on hyperfilm. For western blots of extracellular cyclophilins, extracellular media were collected and centrifuged 5 minutes at 1500g. Thereafter, 200 µl of each supernatant was mixed with 50 µl of 5x sample buffer, and 50 µl of the resulting mix loaded on the western blot.

### Enzymatic activity assays

For determination of alkaline phosphatase (AP), dipeptidyl peptidase-4 (DPP4) and γ-glutamyltransferase (GGT-2) activities, HK-2 cells were grown for 2, 5, and 10 days, and whole cell extracts were prepared with mannitol buffer (50mM D-mannitol, 2mM Tris and 0.1% Triton X-100) supplemented with protease inhibitor cocktail. AP and DPP4 activities were analyzed as described in (71). For AP activity assay, 50μg of protein in a final volume of 50μl of mannitol buffer were incubated with 200 μl of p-nitrophenyl phosphate Liquid Substrate System (#N7653, Sigma-Aldrich) for 15 min at 37 °C and the absorbance was then measured at 405nm. For DPP4 activity assay, 50μg of protein in 90μl of mannitol buffer were mixed with 10 μl of 1.4M glycine-NaOH pH 8.7 and incubated with 100μl of 1.5mM glycyl-L-proline-p-nitroanilide for 30 min at 37 °C. Reaction was stopped by adding trichloroacetic acid. Samples were centrifuged and 50 μl of supernatant were mixed with 50μl of cold 0.2% sodium nitrite and incubated for 10 min at 4 °C. The mixture was then incubated with 50μl of 0.5% ammonium sulfamate for 2 min, at the end of which 100μl of 0.05% n-(1-naphthyl)- ethanediamine were added to the mix, which was further incubated in the dark at 37 °C for additional 30min. Absorbance was then read at 548nm. Finally, GGT activity was analyzed using a Roche/Hitachi Cobas C system. 50 μg of protein in 100 μl of manittol buffer were incubated with L-γ-glutamyl-3-carboxy-4-nitranilide as γ-glutamil donor in the presence of glycyl-glycine and the rate of formation of 5-amino-2-nitrobenzoate was measured spectrophotometrically at 405 nm. All these experiments were performed at least three independent times in triplicate.

### Vectors and Site-directed mutagenesis

For shRNA rescue experiments, wild-type hCypA and wild-type hCypB were cloned into pDONR vectors (pDONR™221, #12536-017, Invitrogen) using the Gateway cloning system (Invitrogen). The QuickChange site-directed mutagenesis kit (Agilent Technologies) was then used to introduce the following silent mutations (c.[151C>A; 153G>A; 156G>T; 159C>T; 162T>C; 165T>C]) in the shRNA targeting sequence of hCypB (see Figure 2A). Since shRNA against hCypA was directed to the 3’ UTR no further modifications were required. The shRNA-rescuing vectors were additionally used to introduce mutations altering PPIase activity of CypA (c.161_162delinsGC; p.Arg55Ala) and CypB (c.259_260delinsGC; p.Arg62Ala). To generate the ER location mutant of hCypB (c.4_75del; p.2_25del), restriction sites were introduced at both sides of the fragment to be deleted. Once all the mutations were performed, all inserts were subcloned to the final destination vector (pLenti CMV Hygro DEST 117-1, Addgene), containing hygromycin resistance, by Gateway recombination. All primers used were generated with the QuikChange Primer Design (Agillent Technologies) tool. Sequences of all constructs were confirmed by DNA sequencing.

### ICC

HK-2 cells were seeded on microscope cover glasses (Marlenfeld GmbH & Co.KG) for 5 days, and, when indicated, starved overnight and treated with 1.5 ng/ml TGFβ for 24 h. Slides were then washed twice in PBS and fixed in 4% paraformaldehyde for 1 h. Aldehyde groups were then blocked with 50 mM NH_4_Cl for 30 min and cells were permeabilized with 0.1% triton X-100 for 10 min. Prior to addition of primary antibodies, unspecific binding sites were blocked with 5% BSA in PBS for 30 min. Slides were then incubated overnight at 4 °C with a 1:100 dilution of primary antibodies and subsequently labeled with secondary fluorescent antibodies (1:200 dilution; see Supplemental Material for references) and Hoechst 33342 (H1399, Invitrogen) for nuclear staining 1 hour at room temperature. Fluorescence labeling was visualized in a confocal spectral FV 1000 Olympus microscope.

### Cell Proliferation

Cell proliferation was measured using carboxyfluorescein diacetate succinimidyl ester (CFSE) staining. Cells were tripsinized, washed with PBS, and incubated with CFSE (Invitrogen) at 2.5 μM final concentration for 10 min at 37 °C in the dark in a cell culture incubator. An aliquot of cells was left unlabeled to set background fluorescence. CFSE was then quenched by washing cells twice with complete medium, and a portion of cells was taken to measure fluorescence at the beginning of the experiment. The rest of labeled cells were plated on six-well plates and incubated at 37°C in 5% CO_2_ for 5 days. Flow cytometric analysis was performed on a FACScalibur (Becton Dickinson) flow cytometer and analyzed with Cell Quest software (Becton Dickenson). Cell proliferation was expressed as the ratio FInd/MFI, where MFI was the median fluorescence intensity of all viable cells at collection and Find the peak fluorescence intensity of the viable non-divided cells.

### TEER and Dextran permeability

For Transepithelial electrical resistance (TEER) and Fluorescein isothiocyanate–dextran (FITC–Dextran) permeability experiments cells were seeded on 24-well transwell plates with 0.4 μm pore polyester membrane inserts (HTS transwell-24 PET; #CLS3379-2EA, Sigma-Aldrich Saint Louis, Missouri, USA) and measurements were performed 5 days after seeding the cells. TEER was measured using an epithelial voltmeter (Millicell-ERS, Millipore, Billerica, MA, USA) with STX100C electrode (for 24 well format) (World Precision Instruments, Sarasota, FL, USA) according to manufacturer’s instructions. For permeability assay, 40 kDa FITC–Dextran (Sigma-Aldrich) in a final concentration of 100 μg/ml was added to the apical compartment of the cells. 180 minutes after adding FITC–Dextran, 200 ul samples were collected from the basolateral compartment and absorbance was measured at 485nm of excitation and 528 nm of emission with a microplate reader (Spectramax Gemini, Molecular Devices, Sunnyvale, CA, USA). Experiments were performed in triplicate with 8 independent samples per group.

### NFκB activity

To determine NFκB activity, 3×10^4^ HK-2 cells were seeded on each well of a 24-well plate and incubated for 24 h. Cells were then cotransfected with 500 ng/well of a DNA mix containing reporter plasmid NFκB (Firefly luciferase gene with NFκB promoter) and reporter plasmid RLTK (Renilla luciferase gene with thymidine kinase promoter) in a 5:1 ratio, plus either negative control plasmid (pCMV-HA) or positive control plasmid (p65 subunit of NFκB). Transfection was performed using Lipofectamine 3000 transfection kit (#L3000-001; Thermo Fisher Scientific, Waltham, MA USA) according to the manufacturer’s instructions. Cells were lysed and luciferase assay was performed using a Dual-Luciferase® Reporter Assay System (#E1910, Promega, Fitchburg, WI, USA). Transfection efficiency was normalized by the value of cotransfected Renilla luciferase.

### Cell Adhesion assay

A cell adhesion assay was performed and modified as described previously (72). HK-2 cells were treated or not with 1,5 ng/ml TGFβ for 48 h. Control and TGFβ-treated HK-2 cells were then trypsinized and washed twice with culture medium to eliminate trypsin. Subsequently, cells were counted and 2×10^4^ cells/well were seeded onto two duplicated 96-well plates. Cells were then allowed to adhere at 37 °C for 30 minutes. After the incubation period, unattached cells from one of the plates were removed by washing twice with PBS. The amount of the remaining attached cells (from the washed plate) and the total of cells (from the unwashed plate) was determined using the XTT assay. For each treatment condition, results were expressed as the ratio of XTT values from washed and unwashed plate.

### Statistics

Results were expressed as the mean ± standard error of the mean (SEM). Experiments performed in mice were analyzed by two-way analysis of variance (ANOVA) followed by Tukey multiple comparison test. For the remaining experiments, Student’s t-test was used for statistical analysis. A P value of less than 0.05 was considered to indicate statistically-significant differences. Statistical analyses were made with commercially available software (GraphPad Prism, version 6.00 for Windows, GraphPad Software, La Jolla California USA).

### Study approval

Animal studies were conducted as approved by the Institutional Animal Care and Use Committee of Mayo Clinic and in accordance with National Institutes of Health guidelines.

## Author Contributions

ES and AM conceived and designed the research studies. ES, MD, AR, AC, KA, MS and JG conducted the experiments. RB provided the CypB KO mice. ES, AM, RB and DB analyzed the data. ES, AM, DB, KA and RB draft the manuscript. All authors read and approved the final manuscript.

## Acknowledgements

Eduard Sarró and Mónica Durán were supported by the generous contribution of Asdent Patients Association. This work was supported in part by grants from Ministerio de Ciencia e Innovación (SAF2011-2950 and SAF2014-59945-R to A. Meseguer), the Fundación Senefro (SEN to A. Meseguer), Instituto de Salud CarlosIII (PIE13/00027), and Red de Investigación Renal REDinREN (12/0021/0013). Karl A. Nath is supported by NIH DK 47060. Meseguer’s research group holds the Quality Mention from the Generalitat de Catalunya (2014 SGR). We want to thank Dr. Santiago Lamas and Dr. José Miguel López-Novoa for critical reading of this manuscript.

## Supplemental Material

**Table S1.** Antibodies (in alphabetical order)

| Antibody | Reference |
| --- | --- |
| Actin | # A5060; Sigma-Aldrich |
| Calnexin | # MA3-027, Thermo Fisher |
| Calreticulin | # 12238; Cell Signaling |
| CD147 | # 376-820 (UM-8D6); Ancell |
| Cyclophilin A | # 07-313; Millipore |
| Cyclophilin B | # PA1-027A; Thermo Scientific |
| E-cadherin | # 610181; BD Transduction Labs |
| ERK1/2 | # 06-182; Merck-Millipore |
| ERK1/2 (Thr202/204) phosphate | # 9101; Cell Signaling |
| Fibronectin | # F6140; Sigma-Aldrich |
| GAPDH | # MA5-15738, Thermo Fisher |
| HA | # 1867423; Roche |
| N-cadherin | # 610920; BD Transduction Labs |
| Occludin | # 71-1500; Invitrogen |
| Pan-Cytokeratin (AE1/AE3) | Dako (Agilent) |
| Slug | # 9585; Cell Signaling |
| Smad2 | # 5339; Cell Signaling |
| Smad2 (Ser465/467) phosphate | # 3108; Cell Signaling |
| Smad3 | # 9523; Cell Signaling |
| Smad3 (Ser423/425) phosphate | # 9520; Cell Signaling |
| Smad4 | # Sc-7966; Santa Cruz Biotechnology |
| Snail | # 3895; Cell Signaling |
| ZO-1 | # 61-7300; Invitrogen |

**Table S2.** Taqman Probes (in alphabetical order)

| Specie | Gene Symbol | Gene Name | Reference |
| --- | --- | --- | --- |
| Human | BMP2 | bone morphogenetic protein 2 | Hs00154192_m1 |
|  | CDH1 | cadherin 1, type 1, E-cadherin (epithelial) | Hs01023894_m1 |
|  | CDH2 | cadherin 2, type 1, N-cadherin (neuronal) | Hs00983056_m1 |
|  | KRT8 | keratin 8 | Hs01670053_m1 |
|  | OCLN | occludin | Hs00170162_m1 |
|  | SKIL | SKI-like oncogene | Hs01045418_m1 |
|  | SMAD6 | SMAD family member 6 | Hs00178579_m1 |
|  | SMAD7 | SMAD family member 7 | Hs00998193_m1 |
|  | SNAI1 | snail family zinc finger 1 | Hs00195591_m1 |
|  | SNAI2 | snail family zinc finger 2 | Hs00950344_m1 |
|  | TWIST1 | twist family bHLH transcription factor 1 | Hs00361186_m1 |
|  | ZEB1 | zinc finger E-box binding homeobox 1 | Hs00611024_m1 |
| Mouse | BMP6 | bone morphogenetic protein 6 | Mm01332882_m1 |
|  | BMP7 | bone morphogenetic protein 7 | Mm00432102_m1 |
|  | BSG | basigin CD147 | Mm0116115_m1 |
|  | CCL2 | chemokine (C-C motif) ligand 2 | Mm 00441242_m1 |
|  | CD68 | CD68 antigen | Mm 03047340_m1 |
|  | CDH1 | cadherin 1 | Mm 01247357_m1 |
|  | COL1A2 | collagen, type I, alpha 2 | Mm00483888_m1 |
|  | FN1 | fibronectin | Mm01256744_m1 |
|  | MMP9 | matrix metalloproteinase 9 | Mm00442991_m1 |
|  | PPIA | peptidylprolyl isomerase A | Mm02342430_g1 |
|  | PPIB | peptidylprolyl isomerase B | Mm00478295_m1 |
|  | SMAD7 | SMAD family member 7 | Mm00484742_m1 |
|  | SNAI1 | snail family zinc finger 1 | Mm00441533_g1 |
|  | SNAI2 | snail family zinc finger 2 | Mm00441531_m1 |
|  | TGFbeta1 | transforming growth factor, beta 1 | Mm01178820_m1 |
|  | TNF | tumor necrosis factor | Mm00443258_m1 |

**Table S3.**
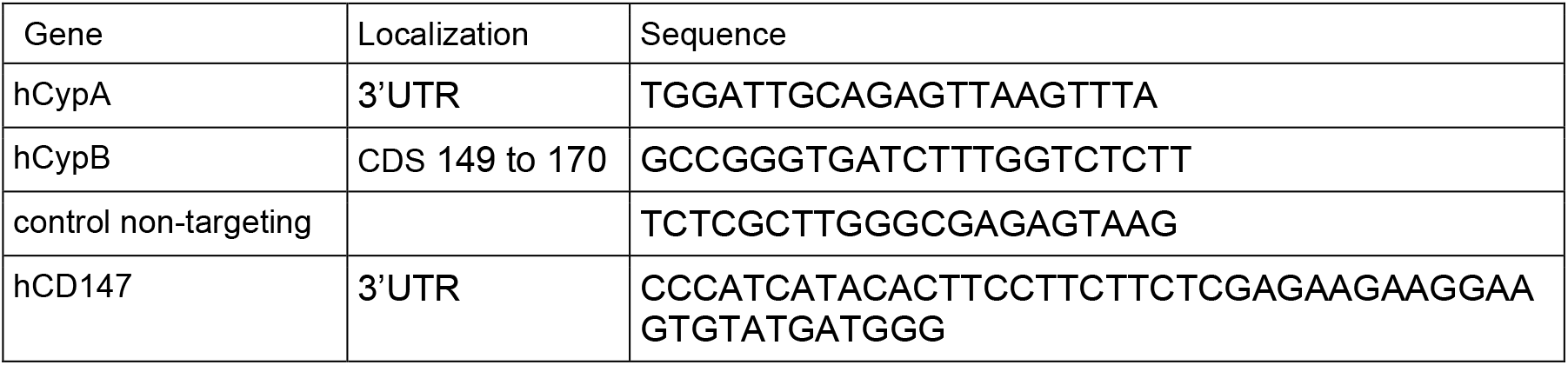
shRNA sequences

**Figure S1.**
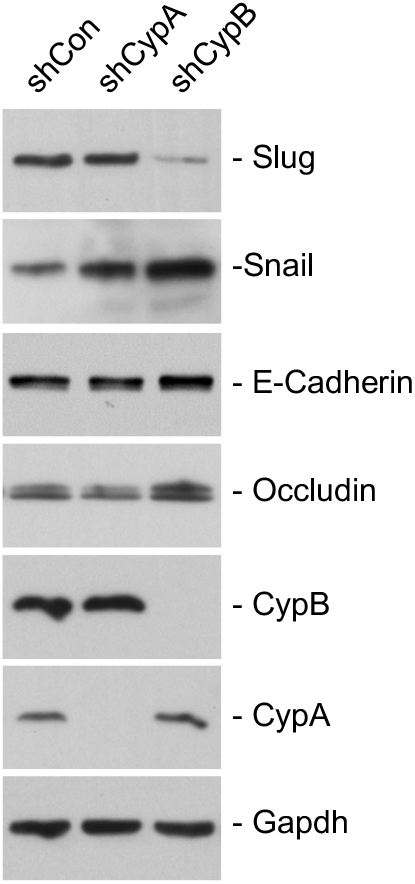
CypB and CypA silencing differentially affects epithelial phenotype of RPTEC. To validate the results observed in HK-2 cells, we silenced CypA and CypB in the human proximal tubule derived cell line RPTEC (#CRL-4031, ATCC). Western Blot shows the decrease in CypA and CypB expression in RPTEC cells stably transfected with shRNA-expressing lentiviral vectors against CypA or CypB, respectively. The expression levels of the epithelial markers E-cadherin and occludin and those of the transcriptional repressors Snail and Slug at 5 days of culture were also analyzed by western blot.

